# Fibroblasts cell lines misclassified as cancer cell lines

**DOI:** 10.1101/166199

**Authors:** Antoine de Weck, Hans Bitter, Audrey Kauffmann

**Affiliations:** Novartis Institutes for Biomedical Research, Basel CH-4002, Switzerland.; Novartis Institutes for Biomedical Research, Cambridge, MA 02139, USA.

## Abstract

Large collections of immortalized cancer cell lines have been widely used as model systems for cancer research and drug discovery. Some of these models however display more fibroblast-like than cancer-like characteristics based on their genetic and genomic characterization. The correct annotation of these cell lines remains a challenge. Here, we report the outcome of our analysis on a large cancer cell line collection, where we found a subset of cell lines misclassified.

---

The present study was undertaken with the aim of curating potential misannotations within cancer cell lines collections. With this goal in mind, we have directed our initial effort at exploring the Cancer Cell Line Encyclopedia (CCLE) (1), which not only represents a large compendium of cancer cell lines commercially available, but also includes cell lines rarely found in other collections, hence often not well characterized. To this end, we have applied a non-linear dimensionality reduction technique, t-distributed Stochastic Neighbour Embedding (t-SNE) (2), to the CCLE gene expression microarray data (Figure 1a). Based on this analysis, the CCLE samples appear mostly clustered according to their respective lineage of origin and, in some instances, even to their sub-lineage as in the case of the hematopoietic samples. Indeed, T-ALL, B-ALL, Hodgkin lymphoma, Non-Hodgkin B cells lymphoma, Non-Hodgkin T cells lymphoma, CML, AML and Multiple Myeloma form distinct clusters. Solid tumor indications display different levels of separation. Some very strongly differentiated, such as melanomas, CRC, non-TNBC and SCLC. Related cancer types from different lineages cluster together, for example ovary and endometrium, stomach and large intestine as well as esophagus and upper aerodigestive tract. The central mixed bulk is formed of outliers from various indications and poorly differentiated types such as the TNBC or NSCLC large cell carcinoma.

**Figure 1.**
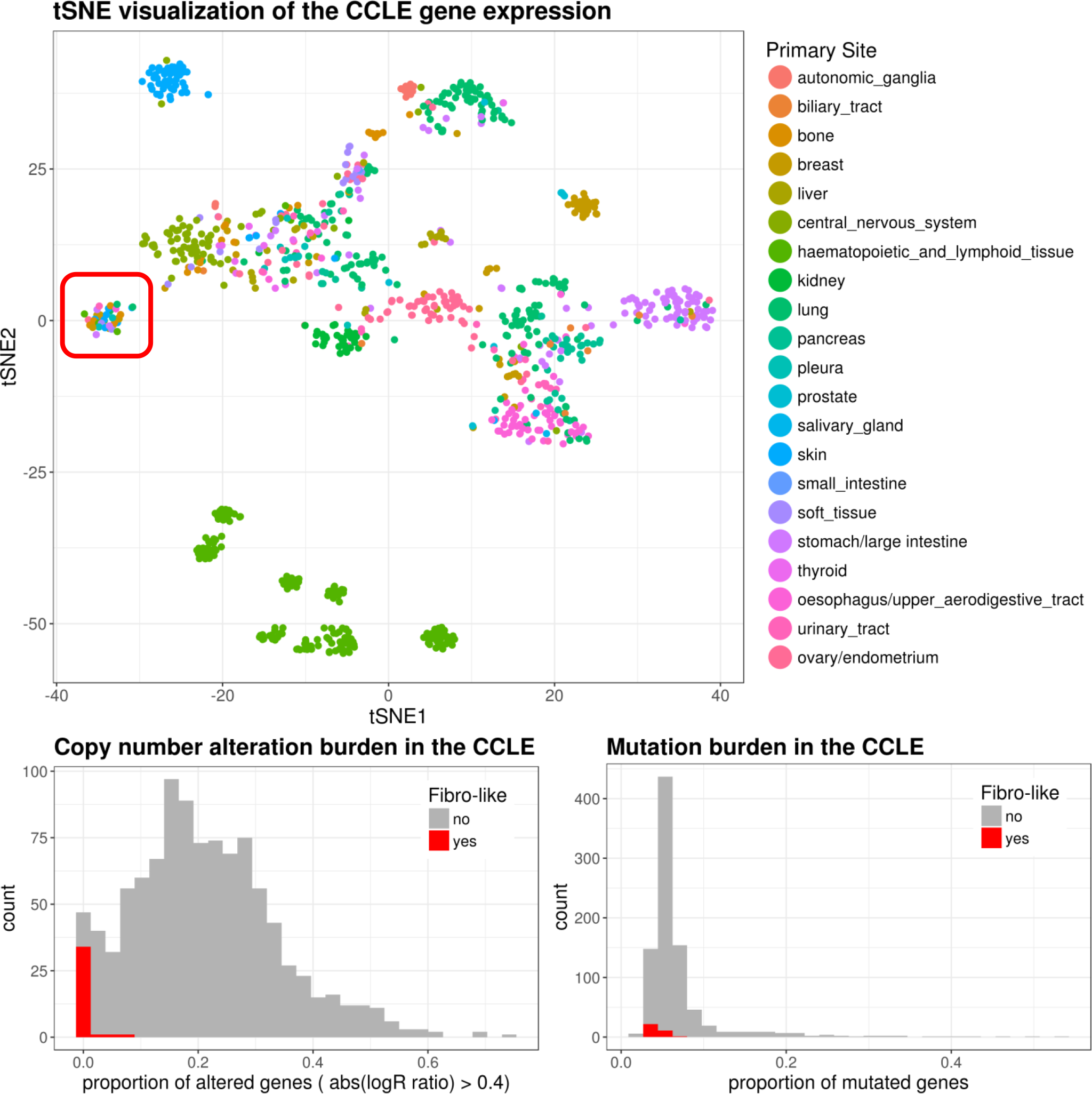
a) tSNE visualization of the CCLE cell lines colored by primary site. The Fibro-cluster is highlighted in red. **b)** Distribution of the proportion of mutated genes across samples. The Fibro-cluster samples are highlighted in red. **c)** Distribution of the proportion of genes with copy number change across the CCLE cell lines. The Fibro-cluster samples highlighted in red.

More interestingly, a population of 40 cell lines forms a clearly segregated cluster that is not defined by its lineage (Table1). Of note, PCA did not yield a similarly distinct cluster (Supplementary figure 1). Application of t-SNE to other CCLE-omics data sets such as RPPA and RNAseq (data not shown) revealed very similar results, confirming that the same cluster of 40 cell lines can be clearly identified by applying t-SNE to gene expression microarray, RNAseq, and to a lesser extent, RPPA data.

Looking into other genomics features of these 40 cell lines, we observed that they exhibit a very low mutation burden in comparison to the overall CCLE population (Figure 1b). Similarly, these lines exhibit a lower burden of copy number alterations in comparison to the rest of the CCLE, as indicated by the low proportion of absolute logR ratio higher than 0.4 (Figure 1c).

Additionally, of those 40 cell lines, 36 were obtained from the American Type Culture Collection (ATCC) for which additional information is available (3). In particular, the morphology annotation was available for 30 of those 36 ATCC lines. Interestingly, 27 of those 30 lines (i.e. 90%) are annotated as having fibroblast morphologies. Outside of this cluster however only 25 out of the 307 annotated ATCC lines display fibroblast morphologies according to the ATCC. This represents a frequency of about 8%, in strong contrast to 90% in the subset identified by this analysis (Supplementary figure 2). Although the morphology annotation was only available for the cell lines of the CCLE which were obtained from the ATCC, the above results strongly suggest that this cluster of 40 cell lines is strongly enriched in cell lines with fibroblast morphologies.

Furthermore, of the 36 ATCC lines in the identified cluster, 35 originated from the Naval Bioscience Laboratory (NBL) collection, for which the ATCC does not guarantee that the morphology and purity will be maintained over time, indicating that those cell lines “may consist of mixtures of stromal and cancer cells in which the former cell type predominates” (4).

Moreover, two lines in the cluster (BJ hTERT and TIG-3 TD) had already been annotated as fibroblasts by the CCLE (5). We also opportunistically included external gene expression data for the known fibroblasts cell lines IMR90 (2 replicates) & WI-38 (3 replicates) in the CCLE dataset (Supplementary figure 3). Interestingly, and despite the expected batch effects due to the lack of common experimental design and normalization, the IMR90 and WI-38 samples cluster closest to our cluster of interest.

All these observations are suggestive of these cell lines having fibroblast-like features, rather than attributes of tumor cells, we are therefore defining this cluster as a “fibro-cluster”.

We also examined potential common features amongst the fibro-cluster. The strongest signal was a strong overrepresentation of translocations/fusions involving COL1A1, COL1A2 and COL3A1, as well as overexpression of these genes (not shown). However we could not find common gene level alterations which could explain the immortality of those lines, suggesting different reprogramming routes, possibly epigenetic.

In order to investigate the existence of fibro-clusters in other large cancer cell lines collections, we applied t-SNE to the Sanger GDSC and the Genentech cancer cell line collections (6,7). As shown in supplementary figures 4 and 5, neither the Sanger nor the Genentech collections seem to exhibit any lines similar to the fibro-cluster.

In this study, we explored the spatial distribution of 3 cancer cell line gene expression datasets by means of t-SNE, leading to the identification of a fibroblast-like cluster of 40 cell lines in one of them (i.e. the CCLE). These cell lines cluster with known fibroblasts expression data, display low CNA and mutation levels and show an overrepresentation in fibroblast morphologies.

Although this population of fibroblasts in the CCLE could at first glance be considered a waste of resource on cell lines of lesser interest, we think those cell lines can provide a very useful and fully characterized control group in the interpretation of contrast experiments. For example for compound profiling, these fibroblast lines could provide a sensitivity baseline for normal-like cells and thus help set an empirical threshold for normal sensitivity.

In summary, we show strong evidence for the presence of an unexpected fibroblast population among the CCLE cell lines which we expect can be beneficially leveraged as a fully characterized control group in the future exploration of this rich dataset.

## Supplementary Information

**Supp. Fig. 1.**
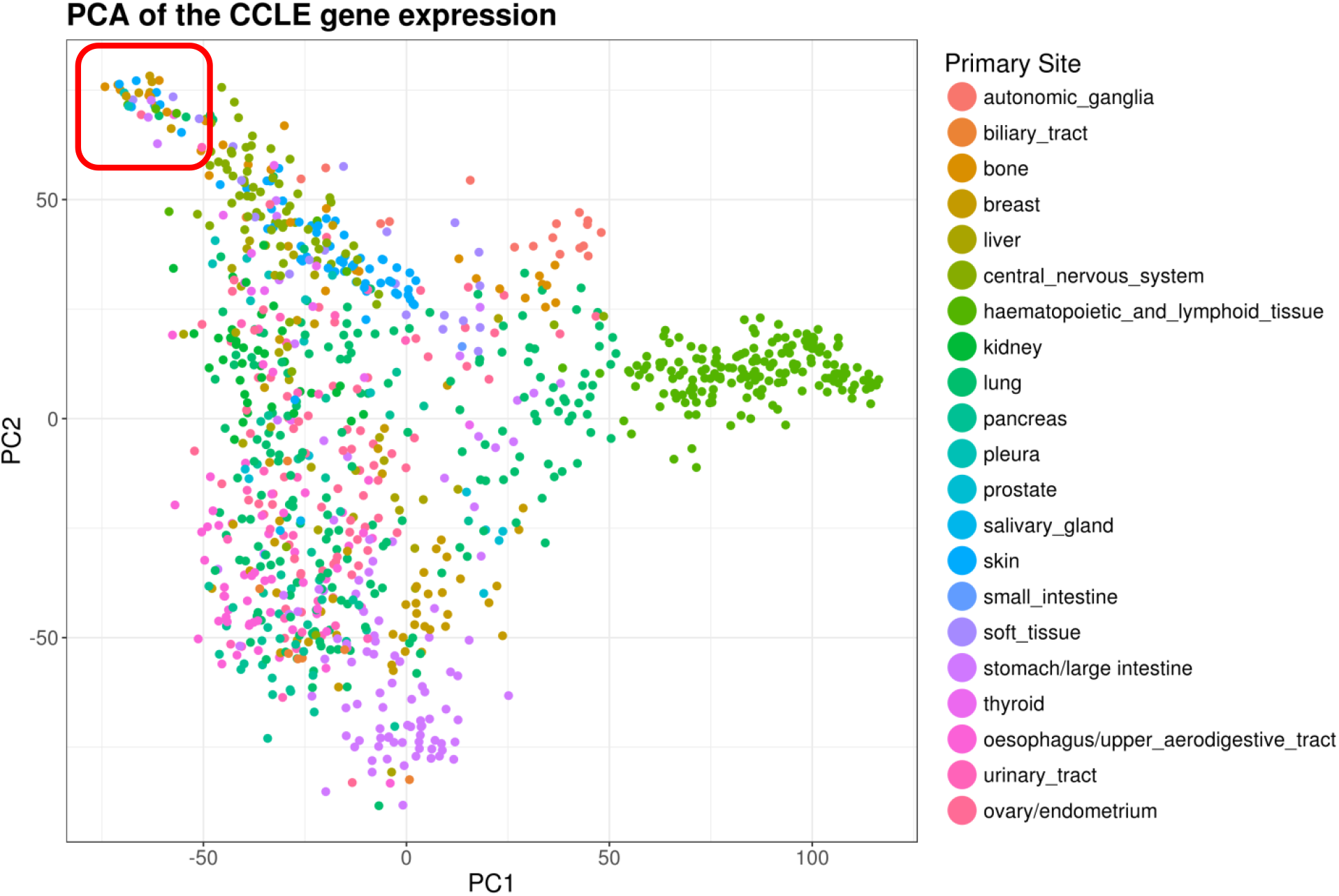
Principle Component Analysis (PCA) of the CCLE expression data.

**Supp. Fig. 2.**
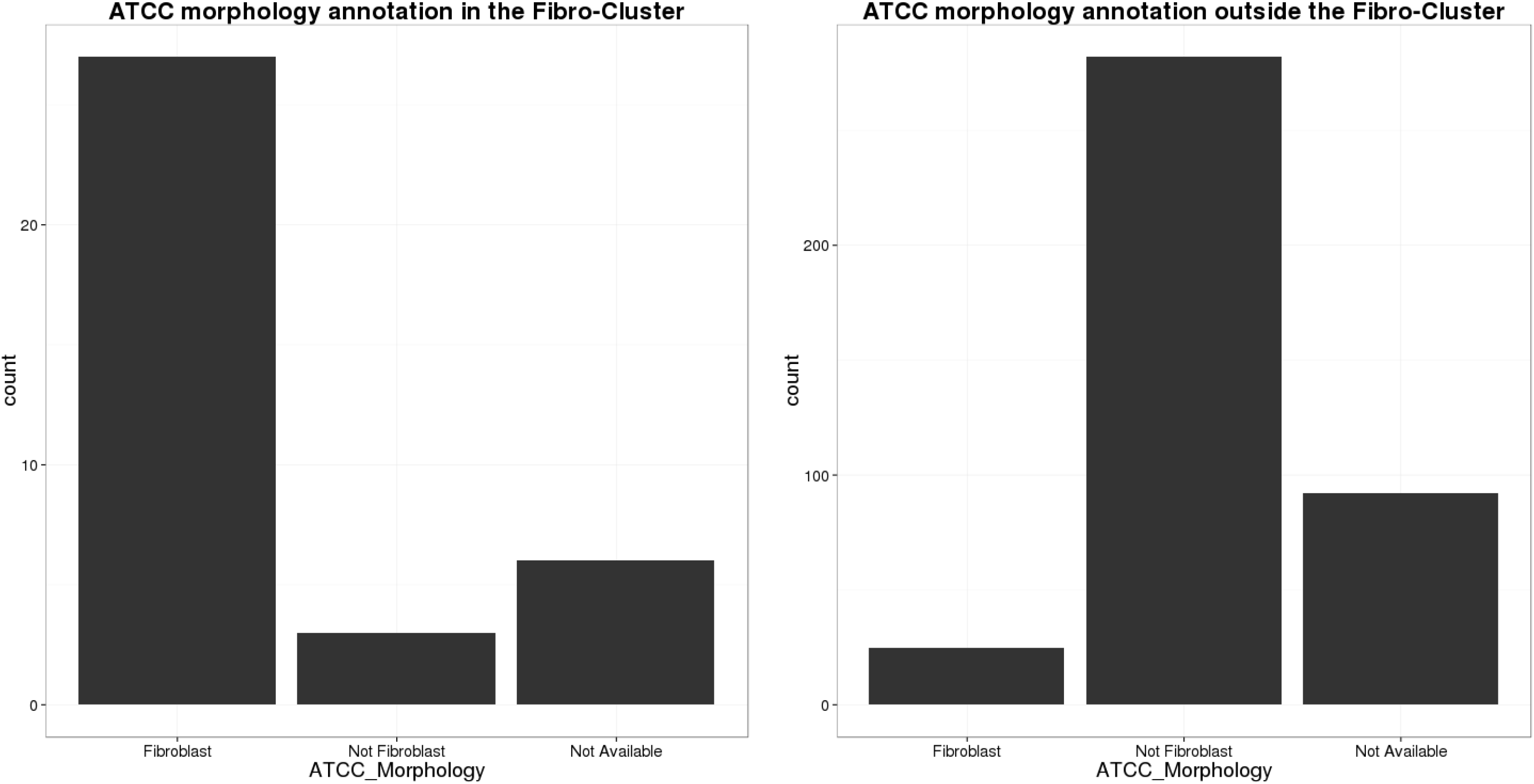
Frequency of ATCC Morphology annotation.

**Supp. Fig. 3.**
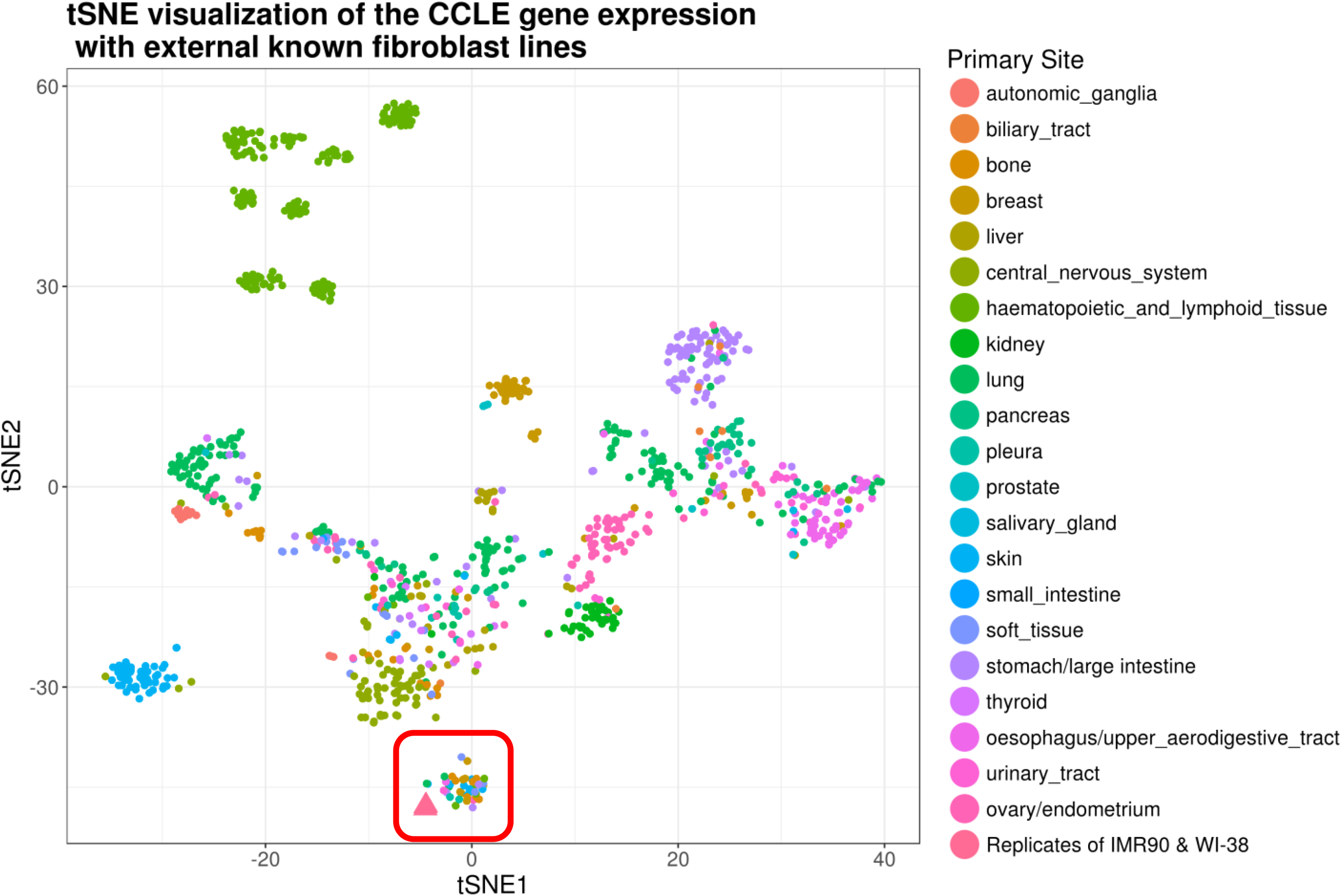
t-SNE of the CCLE expression data including two known fibroblast samples from the literature.

**Supp. Fig. 4.**
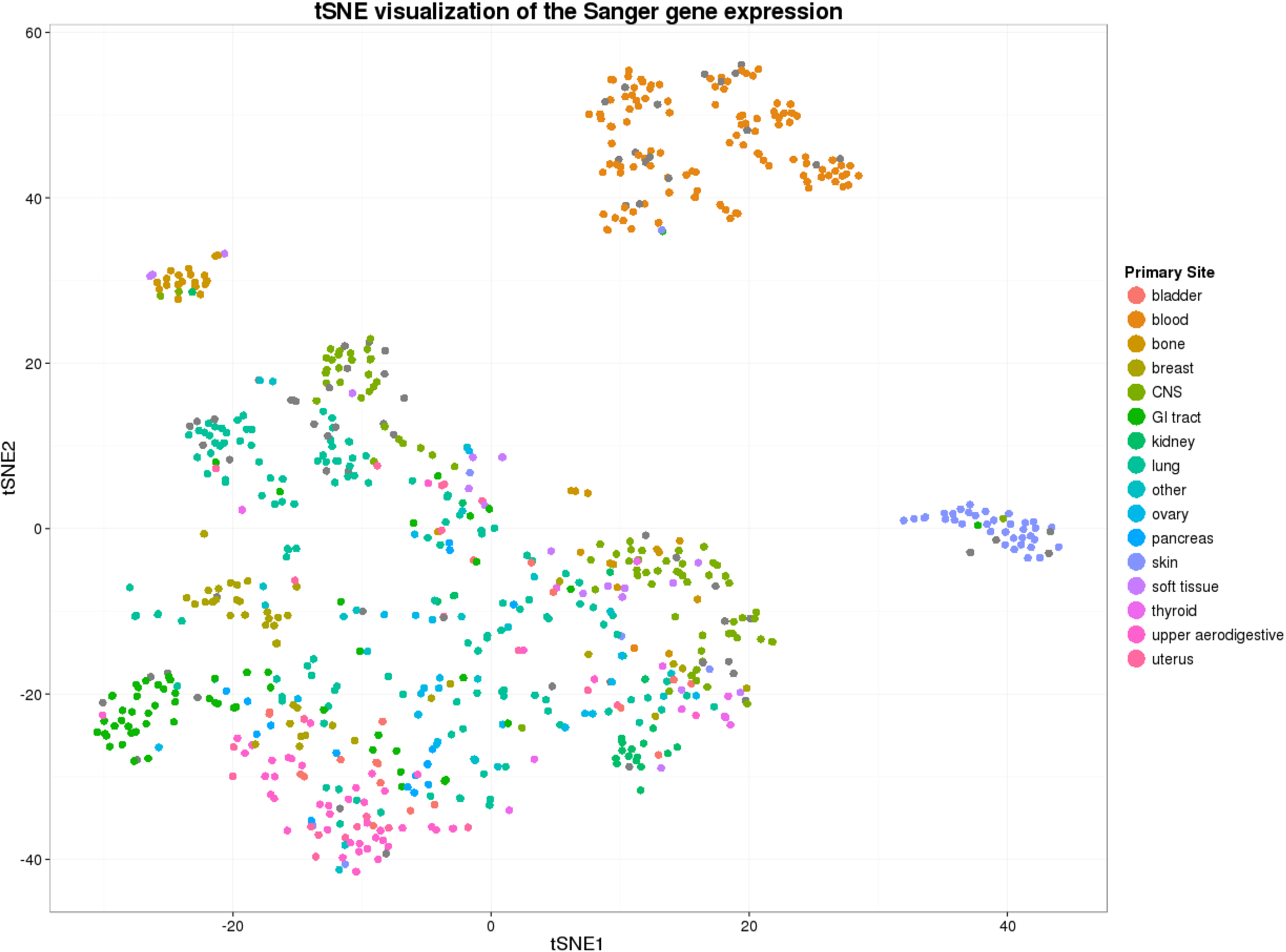
t-SNE of the expression data of the Sanger GDSCC cancer cell line collection.

**Supp. Fig. 5.**
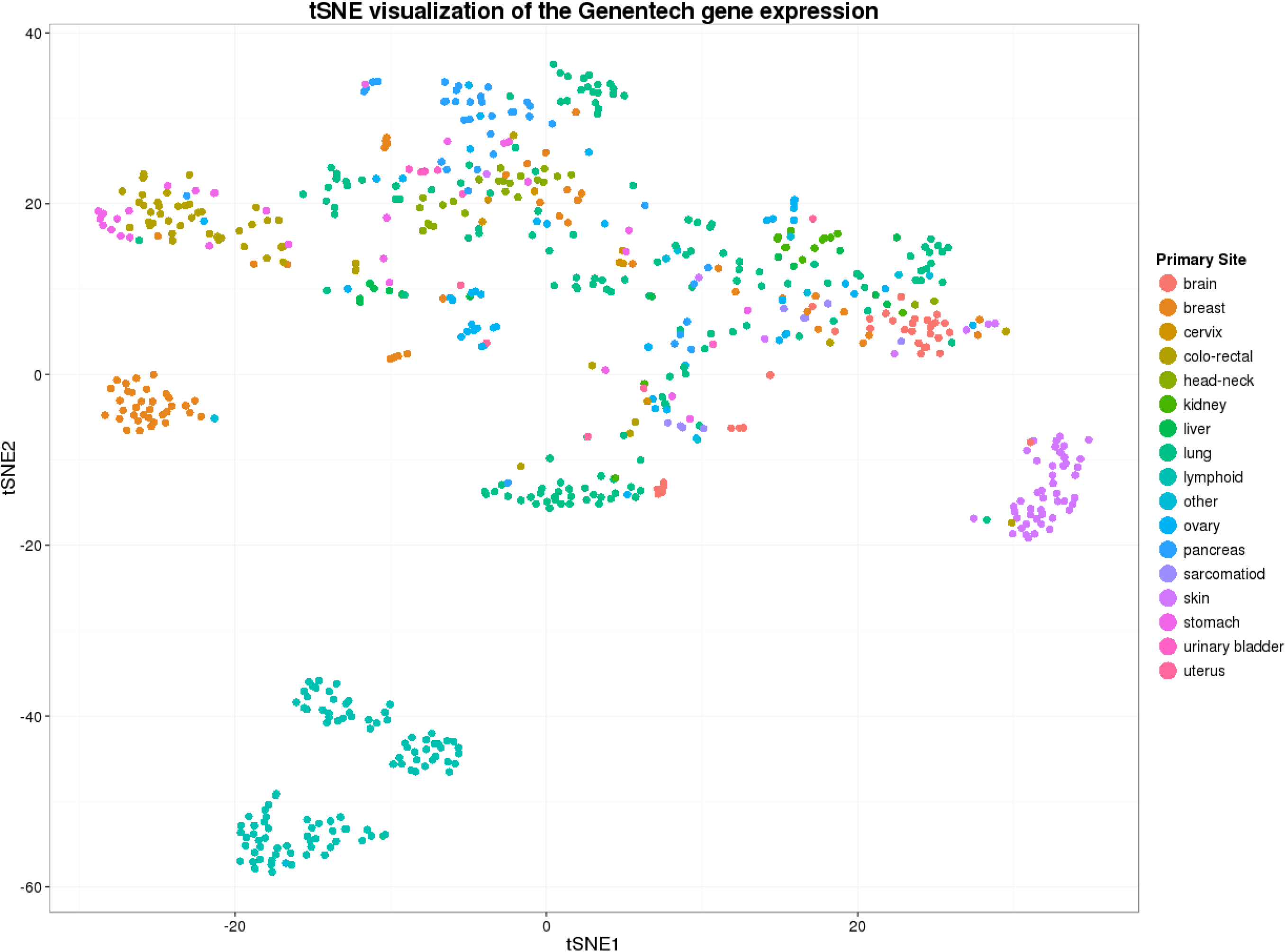
t-SNE of the expression data of the Genentech cancer cell line collection.

**Supp. Table 1.** List of fibro-like cell lines in the CCLE

| CCLE.name | Cell.line.primary.name | Gender | Site.Primary | Histology | Hist.Subtype1 | Source |
| --- | --- | --- | --- | --- | --- | --- |
| HS571T_OVARY | Hs 571.T | F | ovary | carcinoma | NS | ATCC |
| HLFA_LUNG | HLF-a | F | lung | carcinoma | squamous_cell_carcinoma | ATCC |
| HS606T_BREAST | Hs 606.T | F | breast | carcinoma | NS | ATCC |
| HS604T_HAEMATOPOIETIC_AND_LYMPHOID_TISSUE | Hs 604.T | M | haematopoietic_and_lymphoid_tissue | lymphoid_neoplasm | Hodgkin_lymphoma | ATCC |
| HS895T_SKIN | Hs 895.T | F | skin | malignant_melanoma | NS | ATCC |
| HS940T_SKIN | Hs 940.T | M | skin | malignant_melanoma | NS | ATCC |
| HS739T_BREAST | Hs 739.T | F | breast | carcinoma | NS | ATCC |
| T173_BONE | T1-73 |  | bone | osteosarcoma | NS | ATCC |
| HS618T_LUNG | Hs 618.T | F | lung | carcinoma | adenocarcinoma | ATCC |
| HS870T_BONE | Hs 870.T |  | bone | osteosarcoma | NS | ATCC |
| HS675T_LARGE_INTESTINE | Hs 675.T | M | large_intestine | carcinoma | NS | ATCC |
| HS839T_SKIN | Hs 839.T | F | skin | malignant_melanoma | NS | ATCC |
| HS819T_BONE | Hs 819.T | M | bone | chondrosarcoma | NS | ATCC |
| TO175T_HAEMATOPOIETIC_AND_LYMPHOID_TISSUE | TO 175.T | M | haematopoietic_and_lymphoid_tissue | lymphoid_neoplasm | Hodgkin_lymphoma | ATCC |
| HS840T_UPPER_AERODIGESTIVE_TRACT | Hs 840.T | M | upper_aerodigestive_tract | other | papilloma | ATCC |
| HS229T_LUNG | Hs 229.T | M | lung | carcinoma | adenocarcinoma | ATCC |
| HS688AT_SKIN | Hs 688(A).T | M | skin | malignant_melanoma | NS | ATCC |
| HS822T_BONE | Hs 822.T | F | bone | Ewings_sarcoma-peripheral_primitive_neuroectodermal_tumour | NS | ATCC |
| HS737T_BONE | Hs 737.T | F | bone | other | giant_cell_tumour | ATCC |
| HS616T_HAEMATOPOIETIC_AND_LYMPHOID_TISSUE | Hs 616.T | M | haematopoietic_and_lymphoid_tissue | lymphoid_neoplasm | Hodgkin_lymphoma | ATCC |
| HS600T_SKIN | Hs 600.T |  | skin | malignant_melanoma | NS | ATCC |
| TE159T_SOFT_TISSUE | TE 159.T | M | soft_tissue | rhabdomyosarcoma | NS | ATCC |
| HS281T_BREAST | Hs 281.T | F | breast | carcinoma | NS | ATCC |
| HS934T_SKIN | Hs 934.T | F | skin | malignant_melanoma | NS | ATCC |
| HS274T_BREAST | Hs 274.T | F | breast | carcinoma | NS | ATCC |
| HS863T_BONE | Hs 863.T | F | bone | Ewings_sarcoma-peripheral_primitive_neuroectodermal_tumour | NS | ATCC |
| HS821T_BONE | Hs 821.T | F | bone | giant_cell_tumour | NS | ATCC |
| HS343T_BREAST | Hs 343.T | F | breast | carcinoma | NS | ATCC |
| HS888T_BONE | Hs 888.T | M | bone | osteosarcoma | NS | ATCC |
| RS5_PLEURA | RS-5 | M | pleura | mesothelioma | NS | DSMZ |
| HS172T_URINARY_TRACT | Hs 172.T | M | urinary_tract | carcinoma | NS | ATCC |
| HS834T_SKIN | Hs 834.T | F | skin | malignant_melanoma | NS | ATCC |
| TE125T_SOFT_TISSUE | TE 125.T | F | soft_tissue | rhabdomyosarcoma | NS | ATCC |
| DM3_PLEURA | DM-3 | M | pleura | mesothelioma | NS | DSMZ |
| HS698T_LARGE_INTESTINE | Hs 698.T | M | large_intestine | carcinoma | adenocarcinoma | ATCC |
| HS751T_HAEMATOPOIETIC_AND_LYMPHOID_TISSUE | Hs 751.T |  | haematopoietic_and_lymphoid_tissue | lymphoid_neoplasm | Hodgkin_lymphoma | ATCC |
| HS255T_LARGE_INTESTINE | Hs 255.T | F | large_intestine | carcinoma | adenocarcinoma | ATCC |
| HS742T_BREAST | Hs 742.T | F | breast | carcinoma | NS | ATCC |
| BJHTERT_SKIN | BJ hTERT |  | skin | other | immortalized_fibroblast | Academic Lab / Achilles |
| TIG3TD_LUNG | TIG-3 TD |  | lung | other | immortalized_embryonic_fibroblast | Academic Lab / Achilles |

## References

1. Barretina J, Caponigro G, Stransky N, Venkatesan K, Margolin AA, Kim S, et al. The Cancer Cell Line Encyclopedia enables predictive modelling of anticancer drug sensitivity. Nature. 2012 Mar 29;483(7391):603–7.

2. Laurens van der Maaten, Hinton G. Visualizing High-Dimensional Data Using t-SNE. J Mach Learn Res. 2008 Nov;9(Nov):2579–605.

3. ATCC [Internet]. Available from: https://www.lgcstandards-atcc.org/Products/Cells_and_Microorganisms/Cell_Lines/Human/Alphanumeric.aspx

4. The Naval Biosciences Laboratory ATCC Collection [Internet]. Available from: https://www.lgcstandards-atcc.org/Support/Technical_Support/About_NBL_Collection.aspx

5. The Cancer Cell Line Encyclopedia (CCLE) [Internet]. Available from: https://portals.broadinstitute.org/ccle/home

6. Garnett MJ, Edelman EJ, Heidorn SJ, Greenman CD, Dastur A, Lau KW, et al. Systematic identification of genomic markers of drug sensitivity in cancer cells. Nature. 2012 Mar 29;483(7391):570–5.

7. Klijn C, Durinck S, Stawiski EW, Haverty PM, Jiang Z, Liu H, et al. A comprehensive transcriptional portrait of human cancer cell lines. Nat Biotechnol. 2015 Mar;33(3):306–12.

